# Epigenetic-scale comparison of human iPSCs generated by retrovirus, Sendai virus or episomal vectors

**DOI:** 10.1101/355099

**Authors:** Koichiro Nishino, Yoshikazu Arai, Ken Takasawa, Masashi Toyoda, Mayu Yamazaki-Inoue, Tohru Sugawara, Hidenori Akutsu, Ken Nishimura, Manami Ohtaka, Mahito Nakanishi, Akihiro Umezawa

## Abstract

Human induced pluripotent stem cells (iPSCs) are established by introducing several reprogramming factors, such as *OCT3/4*, *SOX2*, *KLF4*, *c-MYC*. Because of their pluripotency and immortality, iPSCs are considered to be a powerful tool for regenerative medicine. To date, iPSCs have been established all over the world by various gene delivery methods. All methods induced high-quality iPSCs, but epigenetic analysis of abnormalities derived from differences in the gene delivery methods has not yet been performed. Here, we generated genetically matched human iPSCs from menstrual blood cells by using three kinds of vectors, i.e., retrovirus, Sendai virus, and episomal vectors, and compared genome-wide DNA methylation profiles among them. Although comparison of aberrant methylation revealed that iPSCs generated by Sendai virus vector have lowest number of aberrant methylation sites among the three vectors, the iPSCs generated by non-integrating methods did not show vector-specific aberrant methylation. However, the differences between the iPSC lines were determined to be the number of random aberrant hyper-methylated regions compared with embryonic stem cells. These random aberrant hyper-methylations might be a cause of the differences in the properties of each of the iPSC lines.

## Introduction

Human induced pluripotent stem cells (iPSCs) are powerful resources for disease modeling, drug discovery, and regenerative medicine because of their potential for pluripotency, ability to self-renew indefinitely, avoidance of rejection of their derivatives by the immune system and for ethical issues [1]. Studies of reprogramming mechanisms and characterization of iPSCs are imperative to ensure the safety of their derivatives in regenerative medicine [2]. Epigenetic reprogramming is an essential event during transformation from somatic cells to iPSCs. DNA methylation is an important epigenetic modification and has a critical role in many aspects of normal development and disease [3–5]. Expression of *OCT-4* and *NANOG* genes, known as reprogramming factors, are induced in restricted tissues with an inverse correlation of DNA methylation during development [6,7]. Transient ectopic expression of defined reprogramming factors forces genome-wide epigenetic exchange and transforms somatic cells to iPSCs [8–12]. After reprogramming, epigenetic profiles of the human iPSCs can be clearly discriminated from the parent somatic cells and are similar to human embryonic stem cells (ESCs), though there is a small fraction of differentially methylated regions [11–15]. In addition, the degree of global DNA methylation in human pluripotent stem cells, ESCs and iPSCs, is higher when compared to somatic cells [11, 12]. This global hyper-methylation of human pluripotent stem cells is a feature shared with primed murine epiblast stem cells, although mouse ESCs, that are naïve state stem cells, have global hypomethylation corresponding to early embryonic cells [16–21].

Human iPSCs have been established by various gene delivery methods. After the first report of iPSC generation by using retrovirus vectors [8], lentivirus vectors [22], Sendai virus vectors [23, 24], PiggyBac vectors [10], plasmid vectors [25], episomal vectors [26], protein transfer [27, 28], mRNA transfer [29], and miRNA transfer [30] have been reported as methods for iPSC generation. All methods induced high-quality iPSCs, but epigenetic abnormalities associated with the specific gene delivery method have not been well analyzed. In this study, we generated human iPSCs derived from menstrual blood cells with three kinds of vectors, i.e., retrovirus, Sendai virus, and episomal vectors, and evaluated them for the scale of genome-wide DNA methylation.

## Results

### Comparison of DNA methylation level in pluripotent stem cells and somatic cells

We generated genetically matched human iPSCs from menstrual blood cells (Edom22) in our laboratory by retrovirus vector infection [12], episomal vector transfection or Sendai viral SeVdp-iPS vector infection (designated as Retro-, Episomal-, and Sendai-iPSCs, respectively) (Fig. 1A and 1B, Supplemental Fig. 1 and 2). To investigate the differences in DNA methylation between iPSCs generated with the three kinds of vectors, we obtained DNA methylation profiles from ESCs, iPSCs including Retro-iPSCs, Sendai-iPSCs, and Episomal-iPSCs, and parent somatic cells, using Illumina’s Infinium HumanMethylation450K BeadChip. Methylation levels are represented as β-values, which range from “0”, for completely unmethylated, to “1”, for completely methylated. Additional data sets from 5 ESCs were obtained from the GEO database [31]. We completely analyzed global DNA methylation of 49 samples (Supplemental Table 1), all with XX karyotypes. All iPSC lines in this study were derived from the same parental somatic cell, Edom22. The promoter regions of pluripotency-associated genes such as *POU5F1*, *NANOG*, *SALL4*, *PTPN6*, *RAB25*, *EPHA1*, *TDGF1* and *LEFTY1* showed low levels of methylation, whereas the promoter regions of somatic cell-associated genes such as *EMILIN1*, *LYST*, *RIN2* and *SP100* were highly methylated in all pluripotent cells (Fig. 1C). These results indicate that all iPSC lines were completely reprogrammed at the core genes, regardless of type of vector used. As assessed by unsupervised hierarchical cluster analysis (HCA) (Fig. 1D) and principal component analysis (PCA) (Fig. 1E) using each iPSC lines (passaged about 30 times), human iPSCs were clearly distinguishable from their parent cells and were similar to ESCs. Sendai-iPSCs appeared to be more similar to ESCs, but no clear difference among the three methods was defined.

**Fig. 1.**
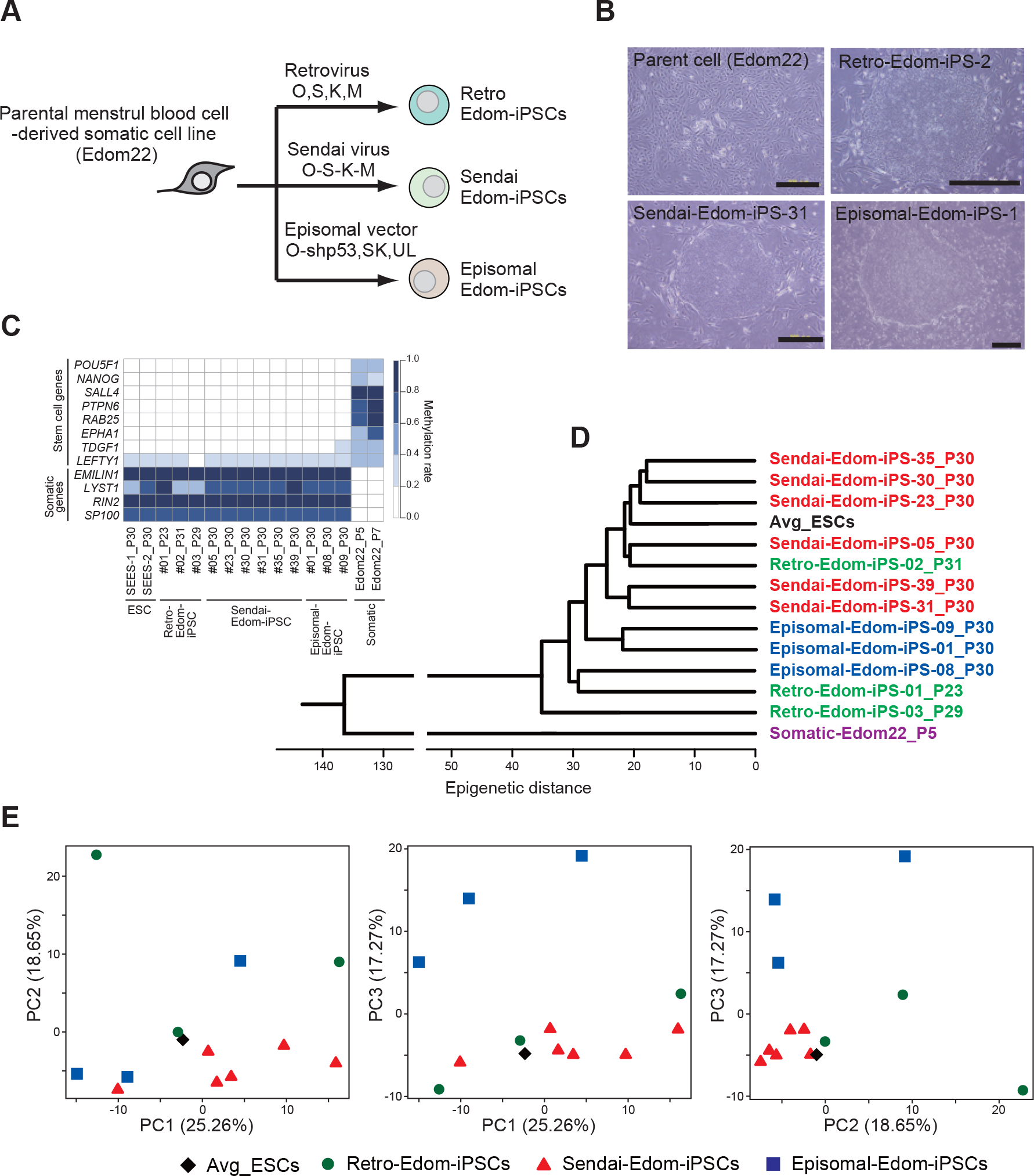
Generation of genetically matched human iPSCs from the same parent somatic cells using three kinds of vetors. A, Schematic for generation of the genetically matched human iPSC lines. Human iPSCs were derived from the same parent somatic cell, Edom-22 using retrovirus, Sendai virus and episomal vectors. B, Morphology of the parent somatic cells and iPSCs. C, Heatmap presenting DNA methylation levels at promoter regions of pluripotency-associated and somatic cell-associated genes. D, Unsupervised hierarchical clustering analysis (HCA) based on DNA methylation scores in the averaged ESCs (black), Retro-Edom-iPSCs (green), Sendai-Edom-iPSCs (red), Episomal-Edom-iPSCs (blue) and the parent comatic cell, Edom-22 (purple). E, Principal component analysis (PCA) based on DNA methylation scores in the averaged ESCs (black diamond), Retro-iPSCs (green circles), Sendai-iPSCs (red triangles) and Episomal-iPSCs (blue squares).

### Identification of differentially methylated regions

In further analysis, we defined a differentially methylated region (DMR) between ESCs and iPSCs as a CpG site whose delta-β-value score differed by at least 0.3 (Fig. 2A). We compared the DNA methylation states of each iPSC line with those of ESCs (the averaged value from 12 ESC lines). The number of DMRs between ESCs and each iPSC line (ES-iPS-DMRs) ranged from 448 to 1,175 in Retro-iPSCs, from 101 to 168 in Sendai-iPSCs, and from 202 to 875 in Episomal-iPSCs. Sendai-iPSCs had lowest number of ES-iPS-DMRs among three kinds of vectors (Fig. 2B). The number of ES-iPS-DMRs in all 6 Sendai-iPSCs was under 168. In Episomal-iPSCs, 2 out of 3 lines showed 300 or fewer ES-iPS-DMRs. Although 1 out of 3 Retro-iPSC lines showed a relatively low number of ES-iPS-DMRs, the range of ES-iPS-DMRs number was wider. These results suggest that the number of ES-iPS-DMRs depended on each cell line rather than on the vectors used for iPSC generation. ES-iPS-DMRs can be categorized into two groups: hyper-methylated and hypo-methylated sites in iPSCs, as compared with ESCs. There was little difference in the number of hypo-methylated ES-iPS-DMRs, especially between Sendai-iPSCs and Episomal-iPSCs (Fig. 2C). The difference between each iPSC line therefore appears to be dependent on the number of abnormal hyper-methylated sites.

**Fig. 2.**
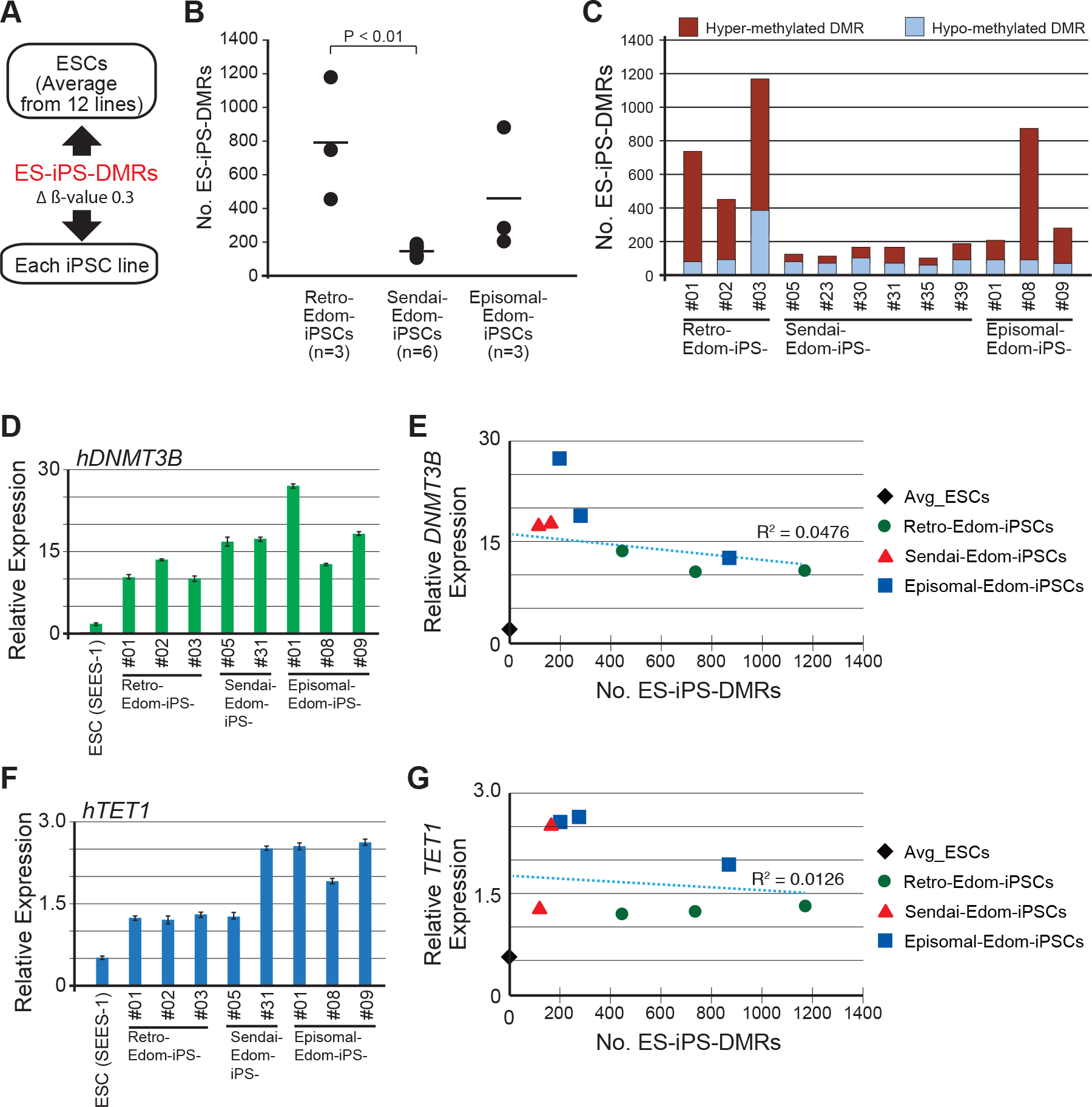
Comparison of differentially methylated regions between ESCs and each iPSC line. A, Comparison of DNA methylation scores for each iPSC lines with that of the average for ESCs. The DMRs between the average for ESCs and iPSCs are designated as ES-iPS-DMRs. B, Comparison of the number of ES-iPS-DMRs in Retro-Edom-iPSCs (n=3), Sendai-Edom-iPSCs (n=6) and Episomal-Edom-iPSCs (n=3). C, Proportion of the hyper- and hypomethylated ES-iPS-DMRs in each iPSC lines. D, Quantitative RT-PCR for hDNMT3B in each PSC lines. E, Correlation between expression level of hDNMT3B and the number of ES-iPS-DMRs. The averaged ESCs, black diamond; Retro-iPSCs, green circles; Sendai-iPSCs, red triangles; Episomal-iPSCs, blue squares. F, Quantitative RT-PCR for hTET1 in each PSC lines. G, Correlation between expression level of hTET1 and the number of ES-iPS-DMRs. The averaged ESCs, black diamond; Retro-iPSCs, green circles; Sendai-iPSCs, red triangles; Episomal-iPSCs, blue squares.

### Correlation between expression of *DNMT*/*TET* genes and ES-iPS-DMRs

In order to investigate whether abnormal hyper-methylation in iPSCs was caused by irregular expression of DNA methyltransferases (DNMTs) and/or Ten-eleven translocations (TETs) that induce de-methylation, we examined expression level of *DNMT* and *TET* genes. All iPSC lines, regardless of type of vector, showed hyper expression of both *DNMT3B* and *TET1* genes compared to ESC (Fig. 2D and 2F). However, there was no correlation between the number of ES-iPS-DMRs and the expression levels of *DNMT3B* or *TET1* gene (Fig. 2E and 2G). Similarly, other *DNMTs* (*DNMT1*, *DNMT3A* and *DNMT3L*) and *TETs* (*TET2* and *TET3*) did not show correlation between the number of ES-iPS-DMRs and expression level.

### Vector-specific ES-iPS-DMRs

We next aimed to detect vector-specific ES-iPS-DMR. We extracted 229 DMRs that overlapped in all 3 Retro-Esom-iPSC lines. The 44 and 76 DMRs also overlapped in all 6 Sendai-Esom-iPSC lines and in all 3 Episomal-Esom-iPSC lines, respectively (Fig. 3A). Approximately 80% of ES-iPS-DMR in each vector group was detected in only one or two lines, suggesting that most of the DMRs occurred randomly in the genome. Comparison of each the overlapping ES-iPS-DMR revealed that 167, 2, 13 ES-iPS-DMRs were detected as the Retro-, Sendai- and Episomal-iPSC-specific DMRs, respectively. Because these aberrant methylation sites at promoter region possibly affect gene expression, we further investigated ES-iPS-DMRs at promoters. While no ES-iPS-DMRs overlapped for all Sendai-iPSC lines, 44 and 5 ES-iPS-DMRs were common in all Retro- and Episomal-iPSC lines at the promoter regions, respectively (Fig. 3B). Details of vector-specific DMRs are summarized in Table 1 and Supplemental Table 2. Interestingly, 5 ES-iPS-DMRs in Episomal-iPSC lines appeared transiently. They were not DMRs at different passages and in different lines (Fig. 3C). In the Retro-iPSC line, 11 out of 44 ES-iPS-DMRs were also transient abnormal regions. These results indicated that Retro-iPSCs had a small subset of the vector-specific ES-iPS-DMRs, but iPSCs generated by non-integrating methods did not have the vector-specific ES-iPS-DMRs, especially those were located promoter regions.

**Fig. 3.**
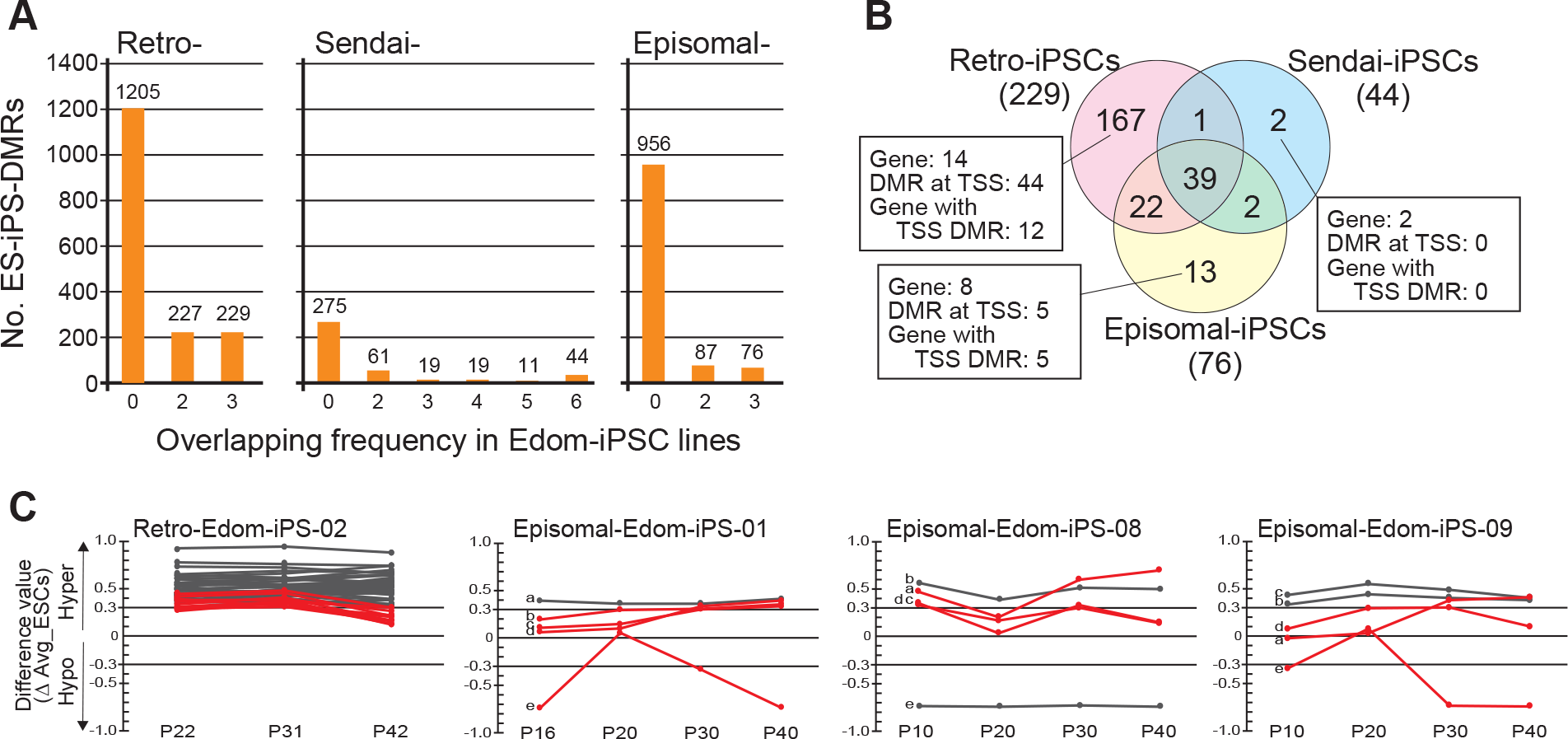
Aberrant methylation in human iPSCs. A, The number of overlapping ES-iPS-DMRs in 3 RetroEdom-iPSC lines, 6 Sendai-iPSC lines and 3 Episomal-iPSC lines. B, Venn diagram showing method-specific DMRs among three kinds of vectors. C, Effect of continuous cultivation on 44 and 5 ES-iPS-DMRs at TSS in Retro-iPSC line and in Episomal-iPSC lines, respectively. The difference values were estimated by subtracting the scores of the averaged ESCs from that of each passage sample. Each line shows a difference value of each probe during culture. Red lines represent transiently ES-iPS-DMRs, which are not DMR at different passage and in different line. a, EPHA10 (probe ID: cg06163371), b, WIPF2 (cg04977733), c, FTH1 (cg25270670), d, ZNF629 (cg05549854), e, SLC19A1 (cg07658590).

**Table 1.** Details of the vector-specific DMRs.

| Type of vector | Retrovirus | Sendai virus | Episomal | Common |
| --- | --- | --- | --- | --- |
| a) No. DMR CpG | 167 | 2 | 13 | 39 |
| b) hyper-methylated DMR in a) | 154 | 0 | 7 | 5 |
| c) hypo-methylated DMR in a) | 13 | 2 | 6 | 34 |
| d) % hyper-methylated DMR (b / a) | 92.22% | 0.00% | 53.85% | 12.82% |
| e) No. DMR CpG on Gene locus in a) | 121 | 2 | 10 | 28 |
| f) % gene locus (e / a) | 72.46% | 100.00% | 76.92% | 71.79% |
| g) hyper-methylated DMR in e) | 117 | 0 | 7 | 2 |
| h) hypo-methylated DMR in e) | 4 | 2 | 3 | 26 |
| i) % hyper-methylated DMR (f / e) | 96.69% | 0.00% | 70.00% | 7.14% |
| Relationship to canonical CpG Island* |  |  |  |  |
| j) CpG island in e) | 94 | 0 | 6 | 4 |
| k) N_Shore in e) | 8 | 1 | 1 | 4 |
| l) N_Shelf in e) | 1 | 0 | 0 | 2 |
| m) S_Shore in e) | 9 | 0 | 2 | 3 |
| n) S_Shelf in e) | 0 | 1 | 0 | 0 |
| o) non-CpG island in e) | 9 | 0 | 1 | 15 |
| Gene region feature category ** |  |  |  |  |
| p) TSS1500 in e) | 42 | 0 | 4 | 4 |
| q) TSS200 in e) | 24 | 0 | 2 | 4 |
| r) 1stExon in e) | 22 | 0 | 2 | 4 |
| s) 5'UTR in e) | 30 | 0 | 3 | 3 |
| t) Body in e) | 94 | 2 | 5 | 60 |
| u) 3'UTR in e) | 2 | 0 | 0 | 9 |
| v) No. Gene in e) | 14 | 2 | 8 | 41 |
| w) No. probe in p) and q) | 44 | 0 | 5 | 6 |
| x) No. Gene with TSS-DMR (v and w) | 12 | 0 | 5 | 6 |

\*Relationship to canonical CpG Island: Shores, - 0-2 kb from CpG island; Shelves, - 2-4 kb from CpG island; N, upstream of CpG island; S, downstream of CpG island.
\*\*Gene region feature category: Gene region feature category describing the CpG position, from UCSC.
TSS200 = 0–200 bases upstream of the transcriptional start site (TSS).
TSS1500 = 200–1500 bases upstream of the TSS.
5'UTR = Within the 5' untranslated region, between the TSS and the ATG start site.
Body = Between the ATG and stop codon; irrespective of the presence of introns, exons, TSS, or promoters.
3'UTR = Between the stop codon and poly A signal.
This information on HumanMethylation450K was provided by Illumina Inc.

### Aberrant methylation in imprinted genes in pluripotent stem cells

We also compared DNA methylation of imprinted genes between somatic cells and iPSCs/ESCs, and identified 413 differentially methylated regions at promoter regions including 68 imprinted genes. These 68 genes comprised 69.4 % of imprinted genes examined. Representative imprinted genes including *MEG3*, *H19*, *PEG3*, *IGFR2*, *PEG10* and *XIST* in ESCs and iPSCs showed aberrant methylation compared with parent somatic cells, independent of the vector type (Fig. 4). In addition, most of the aberrant methylation was not adapted during culture (Supplemental Fig. 3).

**Fig. 4.**
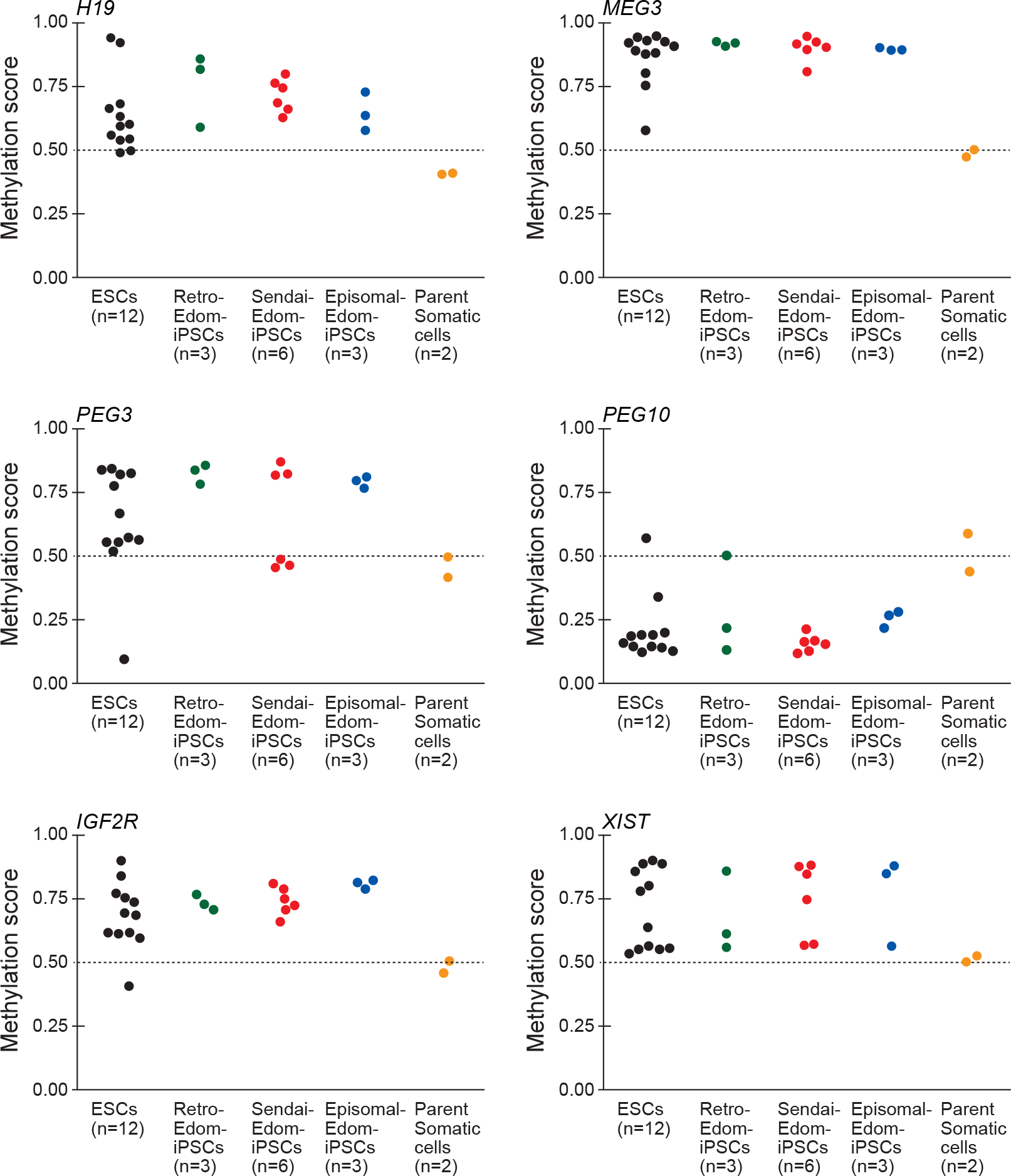
Distribution of methylation scores at promoters of imprint genes and XIST gene. H19, MEG3, PEG3, PEG10, IGF2R and XIST genes are shown. ESCs (n=12, black circles); Retro-Edom-iPSCs (n=3, green circles); Sendai-Edom-iPSCs (n=6, red circles); Episomal-Edom-iPSCs (n=3, blue circles); parent somatic cells, Edom-22 (n=2, yellow circles). The probe IDs and probe positions of each gene were cg17985533, TSS200 for H19, cg09926418, TSS200 for MEG3, cg13960339, TSS200 for PEG3, cg27504782, TSS1500 for PEG10, cg10894318, TSS1500 for IGF2R and cg11717280, TSS200 for XIST. TSS200 and TSS1500 indicated the position of the probe; TSS200, 0 - 200 bases and TSS1500, 200 - 1500 bases upstream of the transcriptional start site (TSS).

## Discussion

Choi et al. [32] reported that hiPSCs, which were generated by using a Sendai virus vector, were molecularly and functionally equivalent to genetically matched hESCs. Schlaeger et al. [33] also reported that there were no substantial method-specific differences in DNA methylation, marker expression levels or patterns, or developmental potential by comparing among Retro-, Lenti-, Sendai-, Episomal-, and mRNA-iPSCs. Our study bridges the gap between the two previous studies. In this study, we compared iPSC lines derived from the same parental somatic cell line, menstrual blood-derived cells, using three types of vectors. Three types of menstrual blood cell-derived-iPSCs are preferable for comparison analysis to avoid any influence from different parental cells. Over 99% of CpG sites in all iPSC lines did not show differences in the methylation levels, compared with ESCs; therefore, iPSCs were almost identical to ESCs in epigenetics regardless of vector types used for iPSC generation. Our findings are consistent with the results of previous studies. However, when focusing the ES-iPS-DMRs of each iPSC line in details, we found that Sendai-iPSCs were more similar to ESCs than Retro- and Episomal-iPSCs at the epigenetic scale. Also, the ranges of the numbers of ES-iPS-DMRs among Retro-iPSC line was wider than those seen in Sendai- and Episomal-iPSCs. This wide variation might result from genome integration of transgenes. Among Episomal-iPSC lines, some lines as well as Sendai-iPSCs showed low numbers of DMRs. The differences between ESCs and iPSCs, especially Sendai- and Episomal-iPSCs, might be derived from characteristics of each of the iPSC lines, rather than type of vectors used for iPSC generation. The type of vector might influence the variety of line-specific properties. The line-specific differences depended on the number of aberrant hyper-methylated DMRs. There was no correlation between this aberrant hyper-methylation and the expression levels of the *DNMT* or *TET* genes; therefore, this aberrant hyper-methylation might be generated at the initial reprogramming step and maintained during culture. Approximately 80% of aberrant hyper-methylated ES-iPS-DMR in each vector group was detected in only one or two lines and there were no vector-specific DMRs in non-integrating methods, suggesting that most of the DMRs occurred randomly in the genome. Therefore, this random hyper-methylation might be a cause of the differences in the properties of each of the iPSC lines.

Aberrant methylation of some imprinted genes in ESCs and iPSCs have been reported by several groups [12, 34, 35]. In this study, we detected 68 imprinted genes that exhibited aberrant methylation, which comprised 69.4% of imprinted genes examined. These abnormalities have been widely detected in pluripotent stem cells. Most iPSC lines as well as ESCs, were abnormally hyper-methylated at *MEG3*, *H19*, *PEG3*, *IGFR2* and *XIST*, regardless of the vector type. The aberrant methylation at imprinted gene and *XIST* gene promoters was maintained throughout continuous passage (data not shown). The aberrant methylation of imprinted genes should therefore be monitored for validation of PSC quality.

In conclusion, we compared genetically matched iPSCs and revealed that there were no vector-specific aberrant methylated regions in iPSCs generated by non-integrating methods. The line-specific properties depended on the number of random aberrant hyper-methylated DMRs rather than type of vectors used for iPSC generation. The differences between the vectors might influence the variety of line-specific properties. It is noteworthy that epigenetic information can be useful to determine iPSC quality.

## Materials and Methods

### Ethics Statement

Human cells were collected with ethical approval of the Institutional Review Board of National Institute for Child Health and Development, Japan. Signed informed consent was obtained from donors, and the specimens were irreversibly de-identified. All experiments handling human cells and tissues were performed in line with the Tenets of the Declaration of Helsinki.

### Human cell culture

Menstrual blood (Edom22) cells were independently established in our laboratory [36, 37], and were maintained in the POWEREDBY10 medium (Glyco Technica Ltd., Sapporo, Japan). Human Retro-iPSCs were generated from Edom22 by retroviral vector pMXs, which encodes the cDNA for human *OCT3/4*, *SOX2*, *c-MYC*, and *KLF4*, via previously described procedures [8] with slight modifications [11, 36, 38, 39]. Episomal-iPSCs were established from Edom 22 by episomal vectors, pCXLE-hOCT3/4-shp53, pCXLE-hSK, and pCXLE-hUL, via procedures described [26]. Sendai-iPSCs were produced from Edom 22 by Sendai viral vector SeVdp-iPS, which encodes the polycisrtonic cDNAs for mouse *Oct3/4*, *Sox2*, *c-Myc*, and *Klf4*, via procedures described [24]. These iPSCs clearly showed human ESC-like characters in terms of morphology; gene expression of stem cell markers;cell-surface antigens; growth (over than 20 passages); normal karyotypes; and teratoma formation (Supplemental Fig. 1 and 2). Non-integrating episomal vectors in the genome or erasing SeVdp vector RNA genome was also confirmed. Human ESCs, SEES, were generated in our laboratory [40]. Human iPSCs and ESCs were maintained on irradiated MEFs in iPSellon medium (Cardio Incorporated, Osaka, Japan) supplemented with 10 ng/ml recombinant human basic fibroblast growth factor (bFGF, Wako Pure Chemical Industries, Ltd., Osaka, Japan). ESC genomes [41, 42] were kindly gifted from Drs. C. Cowan and T. Tenzan (Harvard Stem Cell Institute, Harvard University, Cambridge, MA).

### DNA methylation analysis

DNA methylation analysis was performed using the Illumina infinium assay with the HumanMethylation450K BeadChip (Illumina inc.). Genomic DNA was extracted from the cells using QIAamp DNA Mini Kit (Qiagen). One microgram of genomic DNA from each sample was bisulfite-converted using EZ DNA Methylation kit (Zymo Research), according to the manufacturer’s recommendations. Bisulfite-converted genomic DNA was hybridized to the HumanMethylation450K BeadChip and the BeadChip was scanned on a BeadArray Reader (Illumina inc.), according to the manufacturer’s instructions. Methylated and unmethylated signals were used to compute a β-value, which was a quantitative score of DNA methylation levels, ranging from “0”, for completely unmethylated, to “1”, for completely methylated. On the HumanMethylation450K BeadChip, oligonucleotides for 485,577 CpG sites covering more than 96% of Refseq and 95% of CpG islands were mounted. The probe with MAF (minor allele frequency of the overlapping variant) ≥ 5% [43] and CpG sites with ≥ 0.05 “Detection p value” (computed from the background based on negative controls) were eliminated from the data for further analysis, leaving 416,528 CpGs valid for use with the 49 samples tested. Average methylation was calculated from ESCs, in which 46,681 DMRs among each ESC line in each set were removed. In Fig.1C, each iPSC line (at about the 30th passage) was used and the probes were follows; *POU5F1* (cg15948871, TSS1500), *NANOG* (cg25540142, TSS200), *SALL4* (cg25570495, TSS200), *PTPN6* (cg12690127, TSS200), *RAB25* (cg15896939, TSS200), *EPHA1* (cg02376703, TSS200), *TDGF1* (cg27371741, TSS200), *LEFTY1* (cg15604953, TSS1500), *EMILIN1* (cg19399165, TSS200), *LYST* (cg13677741, TSS1500), *RIN2* (cg17016000, TSS1500), *SP100* (cg23539753, TSS200). TSS200 and TSS1500 indicated the position of the probe; TSS200, 0 - 200 bases and TSS1500, 200 - 1500 bases upstream of the transcriptional start site (TSS).

### Quantitative reversetranscription-PCR

Total RNA was extracted from samples by the conventional method using ISOGEN II (NIPPON GENE, Toyama, Japan). An aliquot of total RNA was reverse-transcribed using ReverTra Ace (TOYOBO, Japan) with random hexamer primers. The cDNA template was amplified using specific primers for *DNMT1*, *DNMT3A*, *DNMT3B*, *DNMT3L*, *TET1*, *TET2* and *TET3*. Expression of glyceraldehyde-3-phosphate dehydrogenase (*GAPDH*) was used as a control. Primers used in this study are summarized in Supplemental Table 3. Quantitative PCR was performed in at least triplicate for each sample on LightCycler®96 Real-Time PCR system (Roche) with Power SYBR Green PCR Master Mix(Life Technologies) using a standard protocol. Relative expression was calculated by the ddCT method using *GAPDH* as an internal standard.

### Accession numbers

NCBI GEO: HumanMethylation450K BeadChip data in this study has been submitted under accession number GSE73938 and GSExxxxxx (in progress). Additional data sets of 5 ESCs were obtained from GSE31848.

## Acknowledgments

We would like to express our sincere thanks to Drs. C. Cowan and T. Tenzan for HUESC lines, to Dr. H. Makino for establishing the Edom22 cells, to Ms. Y. Takahashi for bioinformatics analyses.This research was supported in part by AMED under Grant Number JP18bm0704003 to KoN, and by grants from the Ministry of Education, Culture, Sports, Science, and Technology (MEXT) of Japan; by Ministry of Health, Labor and Welfare (MHLW) Sciences research grant to AU.

---

**Supplemental Fig. 1.**
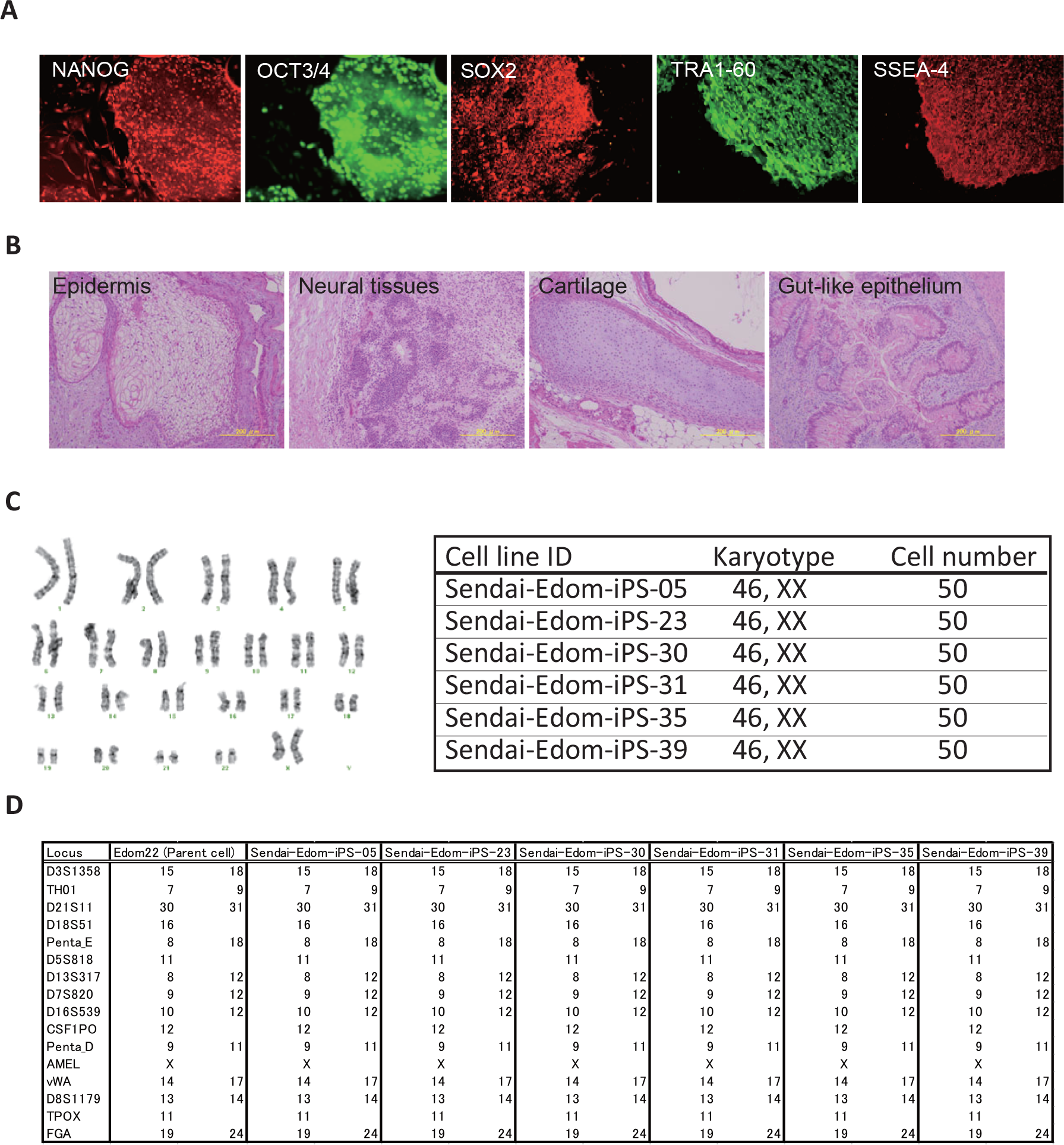
Characterization of Sendai-Edom-iPSCs. A, Immunohistochemistry of stem cell-specific surface antigens, SSEA-4, TRA1-60, and SOX2, OCT3/4, NANOG in Sendai-Edom-iPS-31. B, Teratoma formation of Sendai-Edom-iPS-31 by subcutaneous implantation into NOD/Scid mice. The Sendai-Edom-iPSCs differentiated to various tissues including ectoderm (epidermis and neural tissues), mesoderm (cartilage) and endoderm (gut-like epithelium). Immunostaining and teratoma formation were carried out as previously described [36, 38]. C, Karyotypic analysis of Sendai-Edom-iPSCs. D, STR analysis of Sendai-Edom-iPSCs.

**Supplemental Fig. 2.**
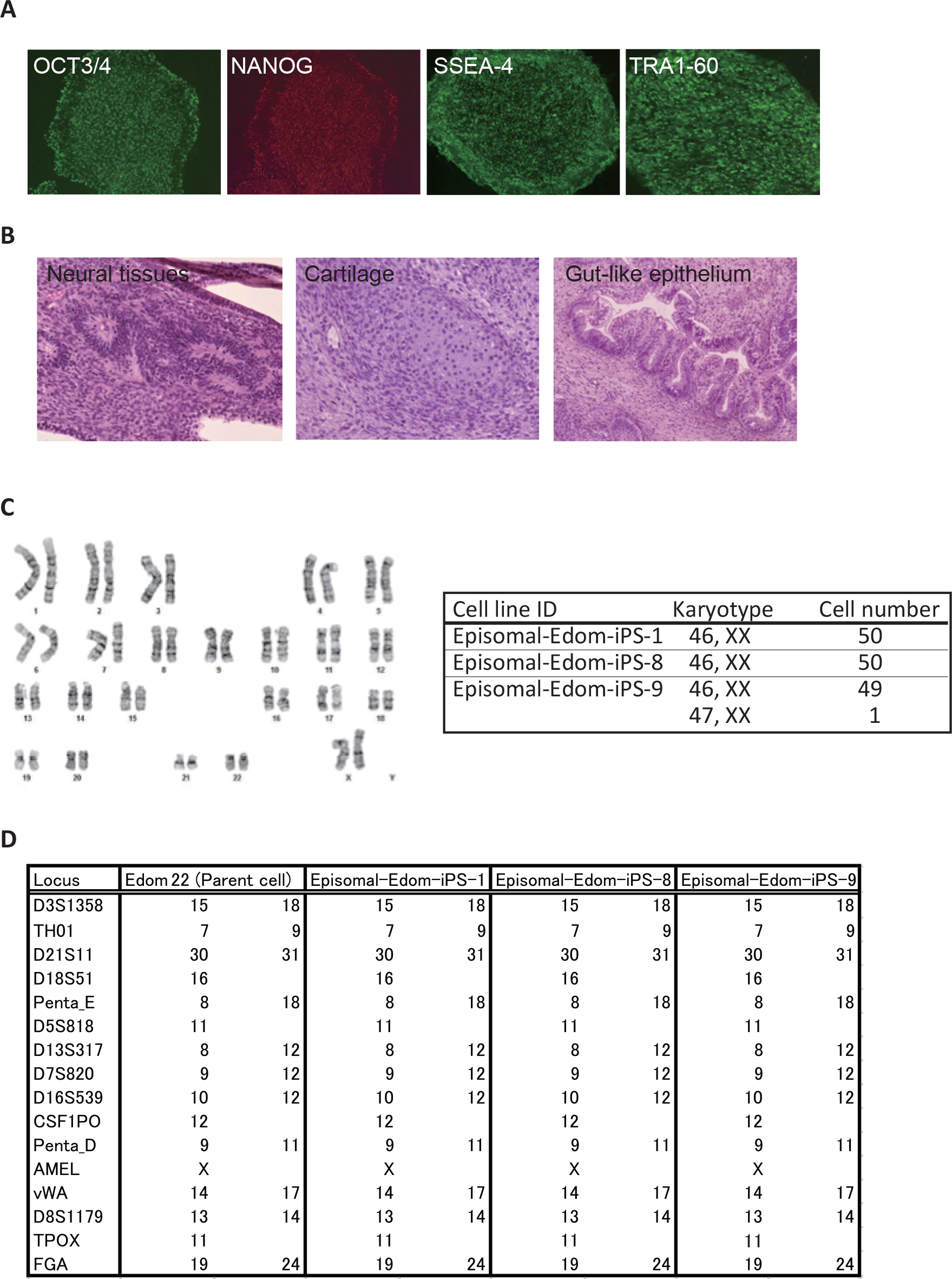
Characterization of Episomal-Edom-iPSCs. A, Immunohistochemistry of stem cell-specific surface antigens, SSEA-4, TRA1-60, and SOX2, OCT3/4, NANOG in Episomal-Edom-iPS-01. B, Teratoma formation of Episomal-Edom-iPS-01 by subcutaneous implantation into NOD/Scid mice. The Episomal-Edom-iPSCs differentiated to various tissues including ectoderm (epidermis and neural tissues), mesoderm (cartilage) and endoderm (gut-like epithelium). Immunostaining and teratoma formation were carried out as previously described [36, 38]. C, Karyotypic analysis of Episomal-Edom-iPSCs. D, STR analysis of Episomal-Edom-iPSCs.

**Supplemental Fig. 3.**
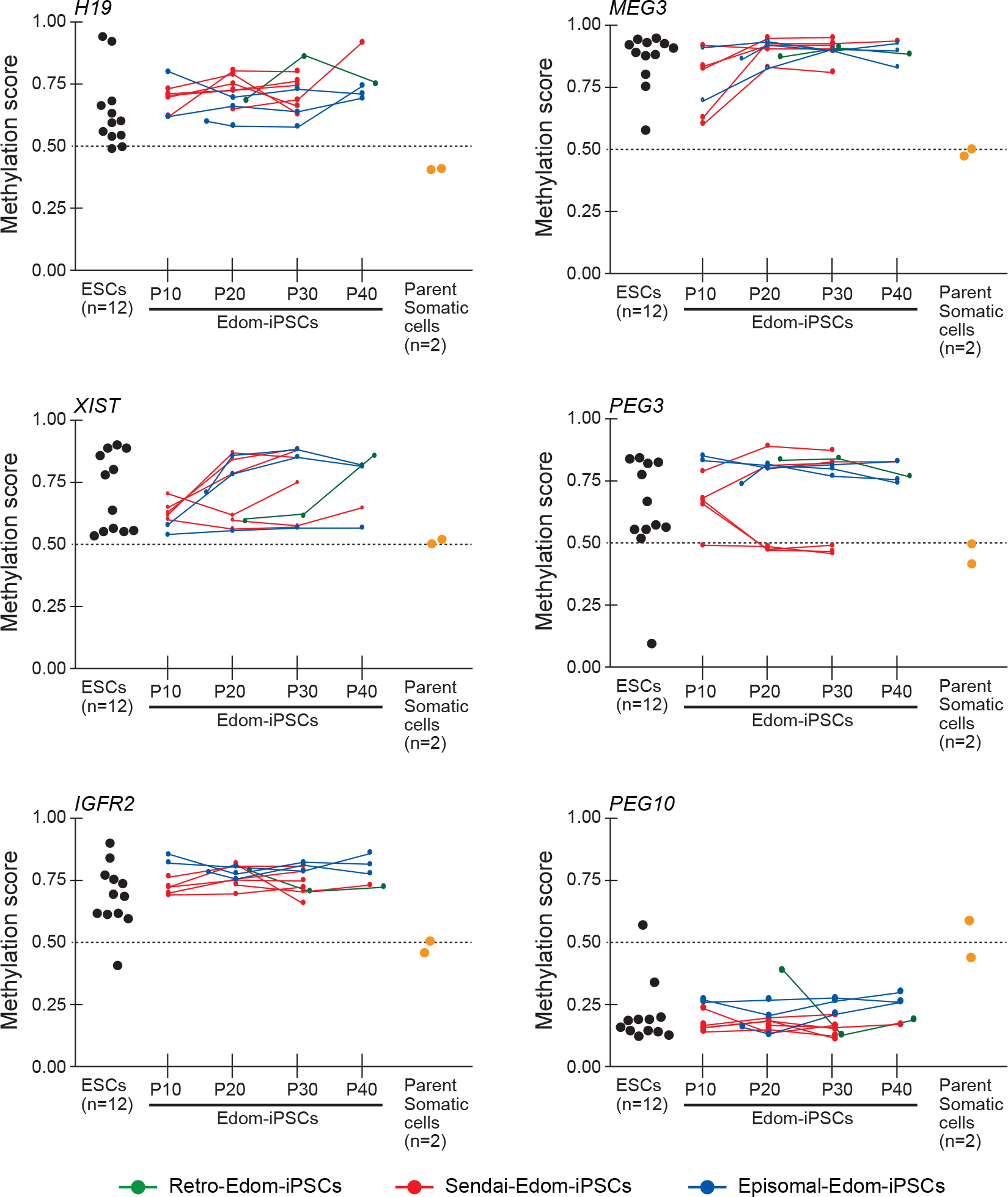
Effect of continuous cultivation on ES-iPS-DMRs at promoters of imprint genes and XIST gene. ESCs (n=12, black circles); Retro-Edom-iPSCs (n=3, green circles and lines); Sendai-Edom-iPSCs (n=6, red circles and lines); Episomal-Edom-iPSCs (n=3, blue circles and lines); parent somatic cells, Edom-22 (n=2, yellow circles). The probe IDs and probe positions of each gene were cg17985533, TSS200 for H19, cg09926418, TSS200 for MEG3, cg13960339, TSS200 for PEG3, cg27504782, TSS1500 for PEG10, cg10894318, TSS1500 for IGF2R and c g11717280, TSS200 for XIST. TSS200 and TSS1500 indicated the position of the probe; TSS200, 0 - 200 bases and TSS1500, 200 - 1500 bases upstream of the transcriptional start site (TSS).

**Supplemental Table 1.**
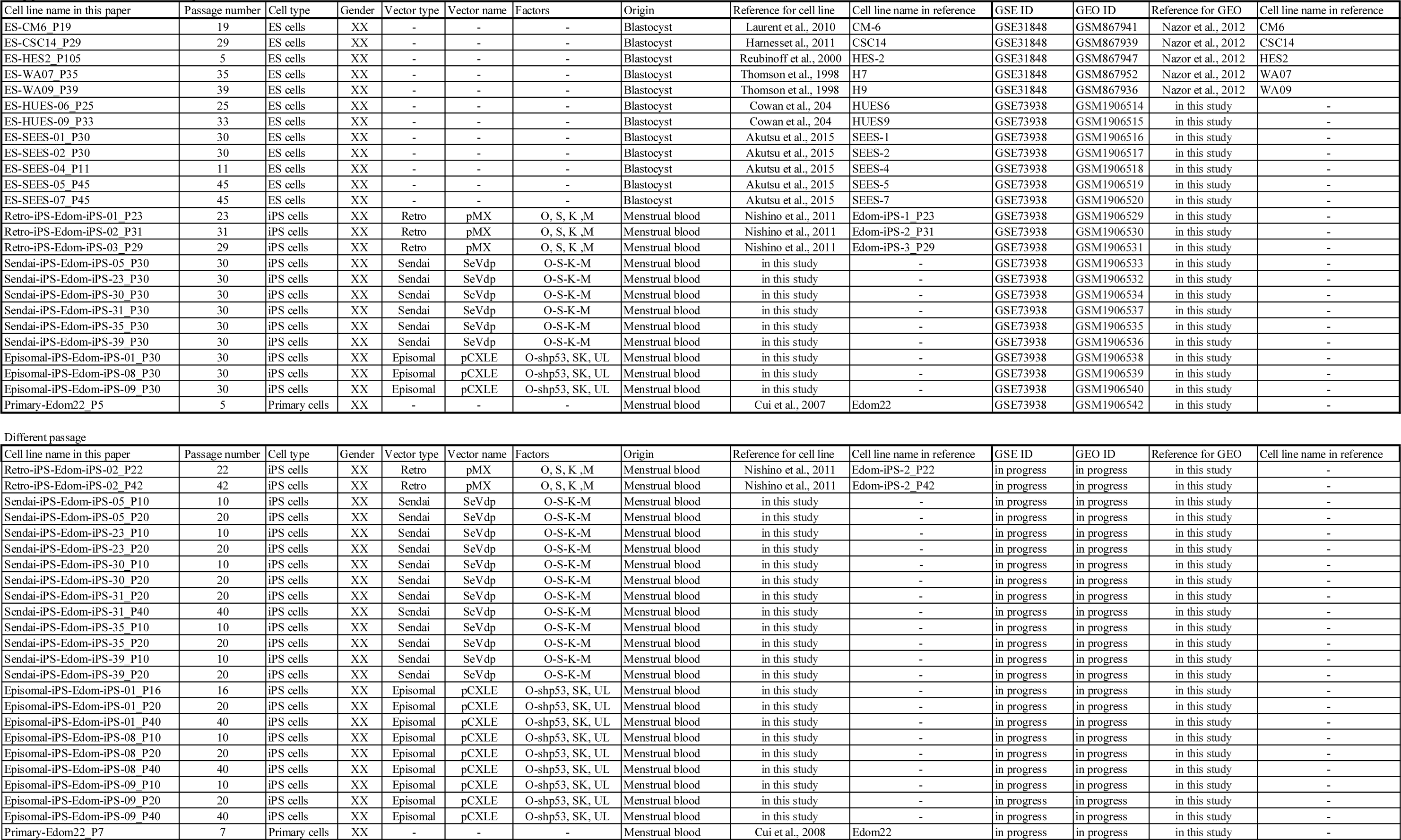
Sample details Different passage

**Supplemental Table 2.** List of gene with DMRs at the promoters Gene with ES-iPS-DMRs at TSS in Retro-Edom-iPSCs Gene with ES-iPS-DMRs at TSS in Episomal-Edom-iPSCs

| UCSC_REFGENE_NAME | description | TargetID |
| --- | --- | --- |
| A2BP1 | Ataxin-2-Binding Protein 1 | cg02230017, cg03586879, cg08545287, cg19378133 |
| CSMD1 | CUB And Sushi Multiple Domains 1 | cg08383399 |
| C5orf38 | Chromosome 5 Open Reading Frame 38 | cg05903444, cg18371475 |
| DPP6 | Dipeptidyl-Peptidase 6 | cg05640128, cg06002683, cg07590961, cg09124223 |
| FZD10 | Frizzled Class Receptor 10 | cg02214745, cg02291403, cg09048530, cg17537177, cg18209212, cg20058043, cg21386611, cg22813950 |
| IRX2 | Iroquois-Class Homeodomain Protein IRX-2 | cg04992127, cg08204280, cg08235864, cg09524455, cg11552694, cg11793269, cg13702053, cg19679633, cg21093166, cg02135861, cg26504021 |
| IRX1 | Iroquois-Class Homeodomain Protein IRX-1 | cg05534710, cg09232937 |
| LOC100190940 |  | cg13822303, cg02681173 |
| OSR1 | Odd-Skipped Related 1 | cg21619325 |
| TCERG1L | Transcription Elongation Regulator 1 Like | cg03943081 |
| TMEM132C | Transmembrane Protein 132C | cg03530754 |
| TMEM132D | Transmembrane Protein 132D | cg00765828, cg12122146, cg13234863, cg23891360, cg25013011 |

Gene with ES-iPS-DMRs at TSS in Episomal-Edom-iPSCs
| UCSC_REFGENE_NAME | description | TargetID |
| --- | --- | --- |
| FTH1 | Ferritin Heavy Chain 1 | cg25270670 |
| WIPF2 | WAS/WASL Interacting Protein Family Member 2 | cg04977733 |
| ZNF629 | Zinc Finger Protein 629 | cg05549854 |
| SLC19A1 | Solute Carrier Family 19 Member 1 | cg07658590 |
| EPHA10 | EPH Receptor A10 | cg06163371 |

**Supplemental Table 4.**
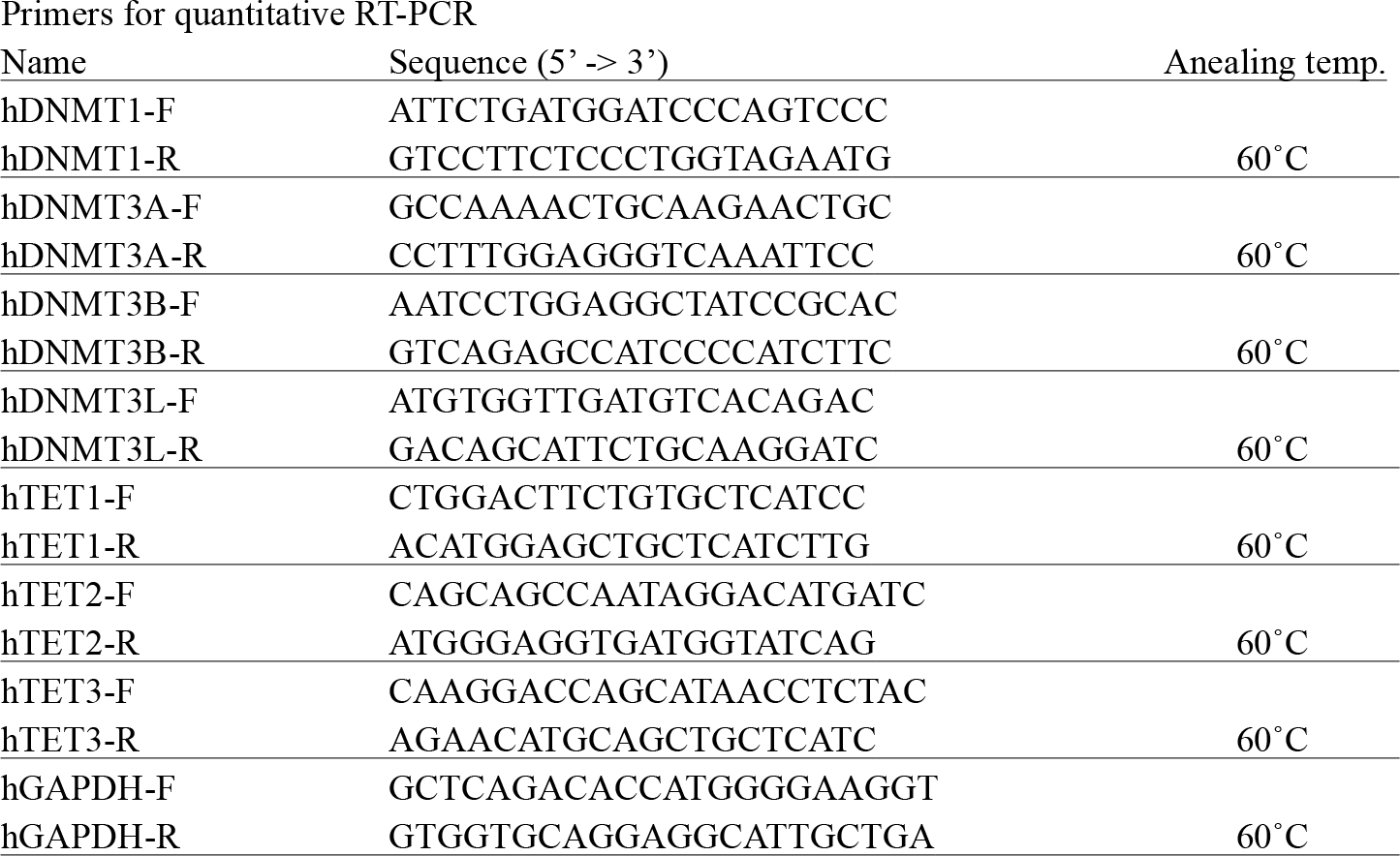
List of primers Primers for quantitative RT-PCR

